# Thick perirenal fat predicts the growth pattern of renal cell carcinoma

**DOI:** 10.1101/606624

**Authors:** Eiji Kashiwagi, Tatsuro Abe, Fumio Kinoshita, Kenjiro Imada, Keisuke Monji, Masaki Shiota, Ario Takeuchi, Junichi Inokuchi, Katsunori Tatsugami, Masatoshi Eto

**Affiliations:** Department of Urology, Graduate School of Medical Sciences, Kyushu University, Fukuoka, Japan

**Keywords:** renal cell carcinoma, visceral fat, perirenal fat, outer location, inner location

## Abstract

**Objective:** To examine the relationship between the direction of renal cell carcinoma growth and the visceral/perirenal fat volume.

Patients and Methods: We retrospectively reviewed computed tomography scans of 153 patients with stage 1 renal cell carcinoma who underwent radical or partial nephrectomy in our hospital between January 2013 and July 2016. We calculated the visceral/subcutaneous/perirenal fat volumes using SYNAPSE VINCENT^®^. Of the 60 patients, the perirenal fat was immunohistochemically stained for leptin, adiponectin, COX-2 and UCP-1, and the association with outward tumor protrusion was evaluated.

**Results:** Of the 153 cases, 88 had confirmed outward expansion (57.5%), 110 were classed as pT1a (52 and 58 with outer and inner expansion, respectively), 43 were classed as pT1b (36 and 7 with outer and inner expansion, respectively; *P*<0.0001). Multivariate logistic regression model showed a trend toward significance in pT1b (vs pT1a, [OR] 6.033, 95%CI=2.409-15.108, P=0.0001), perirenal fat percentage >1.0 (vs ≤1.0, [OR] 2.596, 95%CI=1.205-5.591, P=0.014). as independent predictors for outer protrusion. Immunohistochemical staining was positive for UCP-1 expression in 31 out of 41 outgrowth types (75.6%), and all 19 endogenous types (100%; *P*=0.003).

**Conclusions:** Renal cell carcinoma with thick perirenal fat correlates with an increased likelihood of developing outward tumor protrusion; therefore, fat distribution may affect the development of renal cell carcinoma.

## Introduction

Renal cell carcinoma (RCC) consists of a heterogeneous group of cancers that are derived from the nephron. Various histological and molecular RCC subtypes exist. The international TNM staging system is used to classify RCC because it reflects patient outcomes. The TNM staging system classifies RCC according to the size of the tumor and the grade of extension. In recent years, partial nephrectomy has been the standard surgical treatment for T1a tumors (<4 cm) and select T1b tumors (4–7 cm). Among these tumor classes, the tumor location can be divided into inner or outer. The RCC location is very important to surgeons because intraparenchymal tumors are relatively hard to partially resect and have high perioperative complications [1]. Furthermore, slow-growing RCCs tend to be outwardly located in comparison with rapidly growing types [2]. Therefore, the tumor location may predict surgical complications and malignant potential. However, to date, little attention has been paid to the mechanism of the pattern of RCC growth.

There is lots of epidemiological evidence to suggest that long-standing obesity is one of the primary causes of cancer; including cancers of the breast, colon, esophagus, pancreas and kidney [3]. In the human body there are two types of adipose tissue: white adipose tissue (WAT) and brown adipose tissue (BAT) [4]. WAT is important for energy storage and releases hormones and cytokines that regulate metabolism and insulin sensitivity [5-7]. In contrast, BAT is important for thermogenesis. BAT expresses uncoupling protein 1 (UCP-1), which is rich in mitochondria and uncouples mitochondrial respiration from ATP synthesis to facilitate heat production [5, 8]. BAT is mainly located in thyroid, mediastinal, supraclavicular and perirenal tissues [8]. The kidney is surrounding by perirenal fat; however, no studies have been done to investigate the relationship between RCC and BAT. Perirenal fat is located in the retroperitoneum and is a type of visceral fat [9]. Adipose tissue is generally considered as the storage site of excess energy, although recently is has been revealed that adipocytes have endocrine activity and producing hormones, inflammatory cytokines and adipocytokines [10].

A previous study found that prostate cancer (PCa) patients that had higher periprostatic fat had more aggressive PCa [11]. Moreover, patients with metabolic syndrome, including obesity, have poorer outcomes after radical prostatectomy [12], and obesity is a risk factor for aggressive PCa [13]. For breast cancer patients, obesity is associated with advanced disease at diagnosis and with a poor prognosis [14]. Also in RCC patients, thickness and stranding of perirenal fat affect progression free survival of clinically localized kidney cancer [15]. These results suggest that adipose tissue may also affect RCC growth. In this study, we examined the association between adipose tissue, especially around kidney, and the growth pattern of RCC in patients in our hospital who underwent radical or partial nephrectomy.

## Material and Methods

### Study population

After receiving approval of institutional review board, we retrospectively included 153 patients who were diagnosed with cT1 RCC and who underwent partial or radical nephrectomy. All of the patients were treated at Kyushu University from January 2013 and July 2016. Patients who had hemodialysis were excluded.

### Computed tomography measurement of fat volume

Computed tomography (CT) studies were performed using a 4-slice multidetector CT scanner (Aquilion; Toshiba Medical Systems, Tokyo, Japan). Adipose tissue was identified as the pixels ranging from −250 to −50 Hounsfield units. All imaging data were transferred to a computer workstation for analysis of the total abdominal fat volume. The visceral fat (VF) volume, subcutaneous fat (SF) volume, perirenal fat (PF) volume and abdominal volume were calculated using SYNAPSE VINCENT^®^ software (Fuji Film, Tokyo, Japan). To calculate the visceral fat/subcutaneous fat ratio (V/S ratio), the VF volume was divided by the SF volume. The PF volume was measured by marking the area of adiposity on each CT image (Fig. 1A, B). To calculated the PF percentage, the PF volume was divided by the abdominal volume. To minimize inter-observer variation, adipose tissue assessments were carried out by the same examiner. When the tumor was 50% or more exophytic it was classified as an ‘outer location’ and if it was less than 50% exophytic it was classified as an ‘inner location’.

**Figure 1.**
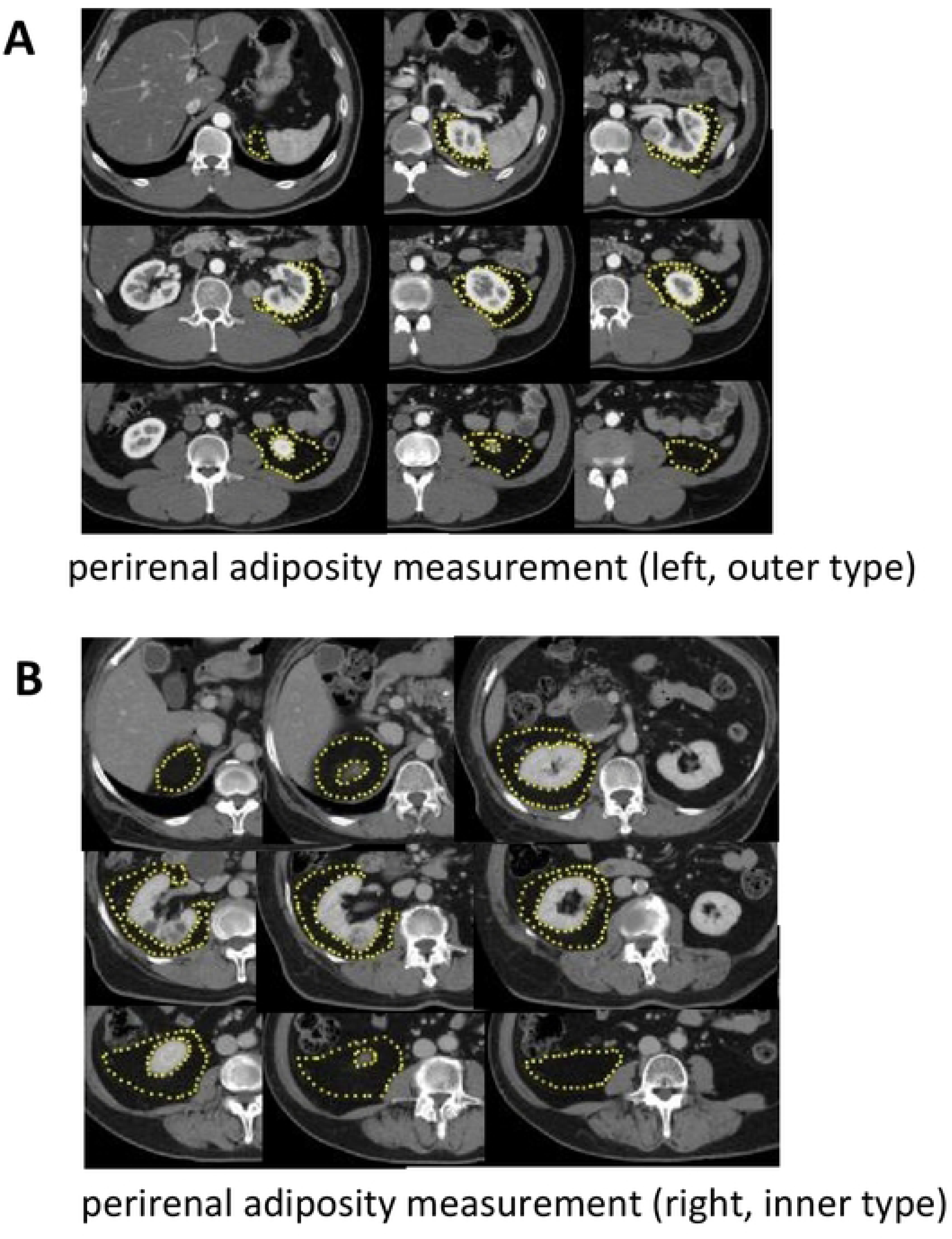
Measurement of the fat area using 5-mm CT slices and SYNAPSE VINCENT^®^ software. (A) Perirenal adiposity measurement (left side, RCC is outer type). (B) Perirenal adiposity measurement (right side, RCC is inner type).

### Immunohistochemistry

Immunohistochemistry was performed on 5-µm thick adipose tissue sections taken from partial or radical nephrectomy specimens which are attached with tumor. Primary antibodies to leptin (dilution 1:100; SC-842, SANTA CRUZ, Dallas, TX, USA), adiponectin (dilution 1:100; ab22554, Abcam, Cambridge, MA, USA), UCP-1 (dilution 1:500; U6382, Sigma, St Louis, MO, USA), or COX-2 (dilution 1:100; 160112, Cayman, Ann Arbor, MI, USA) were applied, followed by a broad-spectrum secondary antibody (Invitrogen, Carlsbad, CA, USA), as described previously [16].

For scoring leptin, adiponectin, UCP-1 and COX-2 expression, stained cells were divided into three categories as follows: 0, negative; 1, positive; and 2, strong positive. All stains were visually quantified by a single pathologist (T. A.) blinded to the sample identity.

### Statistical analysis

All statistical analyses were performed using JMP13 software (SAS Institute, Cary, NC, USA). Univariate and multivariate analyses were performed using the logistic regression model. Correlations between parameters were examined by χ^2^ test. *P* values <0.05 were considered as significant.

## Results

### Clinical characteristics

The clinical features of the patients are shown in Table 1. The median age at diagnosis was 62 (34–83) years, and of the 153 patients, 112 (73.2%) were men and 41 (26.8%) were women. Seventy-three patients (47.7%) had right side RCC and 80 (52.3%) had left side RCC; 88 (57.5%) were outer and 65 (42.5%) were inner expansion; 132 (86.3%) had clear cell RCC, 11 (7.2%) had papillary RCC and 9 (5.9%) had chromophobe RCC; 110 (71.9%) of the patients presented with pT1a tumors and 43 (28.1%) with pT1b; 23 (15.0%) of the patients had Fuhrman nuclear grade 1, 105 (68.6%) had grade 2, 24 (15.7%) had grade 3 and 1 (0.7%) had grade 4. The median body mass index (BMI) was 23.5 (16.0–36.5). The median V/S ratio was 1.00 (0.11–2.73) and the median perirenal fat percentage was 1.01 (0.03–3.59).

**Table 1.** Patient characteristics.

| variable |  |
| --- | --- |
| Median age, Years (range) | 62 (34-83) |
| Gender, n (%) |  |
| male | 112 (73.2%) |
| female | 41 (26.8%) |
| Laterarity, n (%) |  |
| right | 73 (47.7%) |
| left | 80 (52.3%) |
| Expansion pattern, n(%) |  |
| outer | 88 (57.5%) |
| inner | 65 (42.5%) |
| Histology, n (%) |  |
| clear cell | 132 (86.3%) |
| papillary | 11 (7.2%) |
| chromophobe | 9 (5.9%) |
| collecting duct | 1 (0.6%) |
| pT stage, n (%) |  |
| pT1a | 110 (71.9%) |
| pT1b | 43 (28.1%) |
| Fuhrman nuclear grade, n (%) |  |
| 1 | 23 (15.0%) |
| 2 | 105 (68.6%) |
| 3 | 24 (15.7%) |
| 4 | 1(0.7%) |
| Median BMI, kg/m<sup>2</sup> (range) | 23.5 (16.0-36.5) |
| Median visceral fat, % (range) | 33.0 (5.1-59.9) |
| Median subcutaneous fat, % (range) | 15.7 (2.0-37.9) |
| Median body fat, % (range) | 32.8 (5.0-54.8) |
| Median V/S ratio, ratio (range) | 1.00 (0.11-2.73) |
| Median Perirenal fat, % (range) | 1.01 (0.03-3.59) |
| Hypertension, n (%) |  |
| Negative | 76 (49.7%) |
| Positive | 77 (50.3%) |
| Diabetes mellitus, n (%) |  |
| Negative | 137 (89.5%) |
| Positive | 16 (10.5%) |
| Hyperlipidemia, n (%) |  |
| Negative | 121 (79.1%) |
| Positive | 32 (20.9%) |

V/S ratio: Visceral fat/ subcutaneous fat ratio
Body fat percentage: (Visceral fat + Subcutaneous fat)/abdominal volume
Perirenal fat percentage: perirenal fat/ abdominal volume

### Univariate and multivariate analyses between characteristics of patients and expansion pattern

Table S1 summarizes the expansion pattern and characteristics of the patients. To determine what are the important factors to grow outward, we performed univariate and multivariate analyses. In the univariate analysis, gender (odds ratio [OR] 2.129, 95% confidence interval (CI) 1.030-4.400, P=0.041), pT stage ([OR] 5.736, 95% CI 2.351-13.995, *P*=0.0001) and PF percentage divided by 1.0 ([OR] 2.713, 95% CI 1.399-5.258, *P*=0.003) were associated with expansion pattern (Table 2). In the multivariate analyses showed that the pT stage ([OR] 6.033, 95% CI 2.409-15.108, *P*=0.0001) and PF percentage ([OR] 2.596, 95% CI 1.205-5.591, *P*=0.014) are independent factors.

**Table 2.**
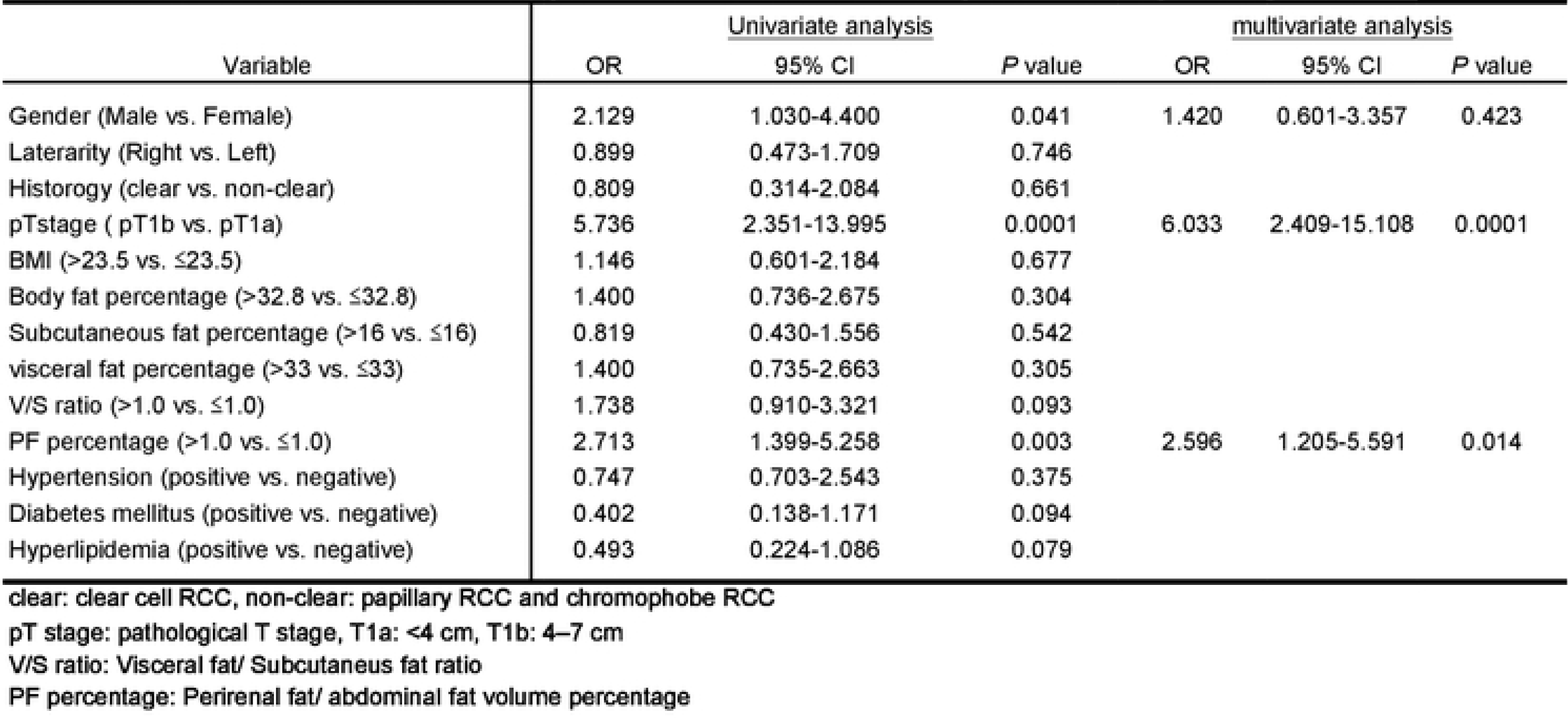
Univariate and multivariate analyses between characteristics of patients and expansion pattern.

Leptin, adiponectin, COX-2 and UCP-1 expression in the perirenal fat of RCC patients We immunohistochemically stained for leptin, adiponectin, COX-2 and UCP-1 in samples from the 60 patients. Positive signals representing these proteins were predominantly detected in adipocytes (Fig. 2), and their expression patterns are summarized in Table 3. Leptin was detected in 3 (7.4%) of the 41 outer expansion samples (3 [7.4%] 1+) and 2 (10.5%) of the 19 inner expansion samples (2 [10.5] 1+; *P*=0.681, 0 vs 1+/2+). Adiponectin was positive in all 41 (100%) of the outer expansion samples (3 [7.3%] 1+, 38 [92.7%] 2+) and all 19 (100%) of the inner expansion samples (1 [5.2%] 1+, 18 [94.8%] 2+; *P*=0.762, 0/1+ vs 2+). COX-2 was also positive in all 41 (100%) of the outer expansion samples (41 [100%] 2+) and all 19 (100%) of the inner expansion samples (1 [5.2%] 1+, 18 [94.8%] 2+; *P*=0.126, 0/1+ vs 2+). UCP-1 was positive in 31 (75.6%) of the 41 outer expansion samples (31 [75.6%] 1+) and all 19 (100%) of the inner expansion samples (14 [73.7%] 1+, 5 [26.3%] 2+; *P*=0.003, 0 vs 1+/2+).

**Table 3.**
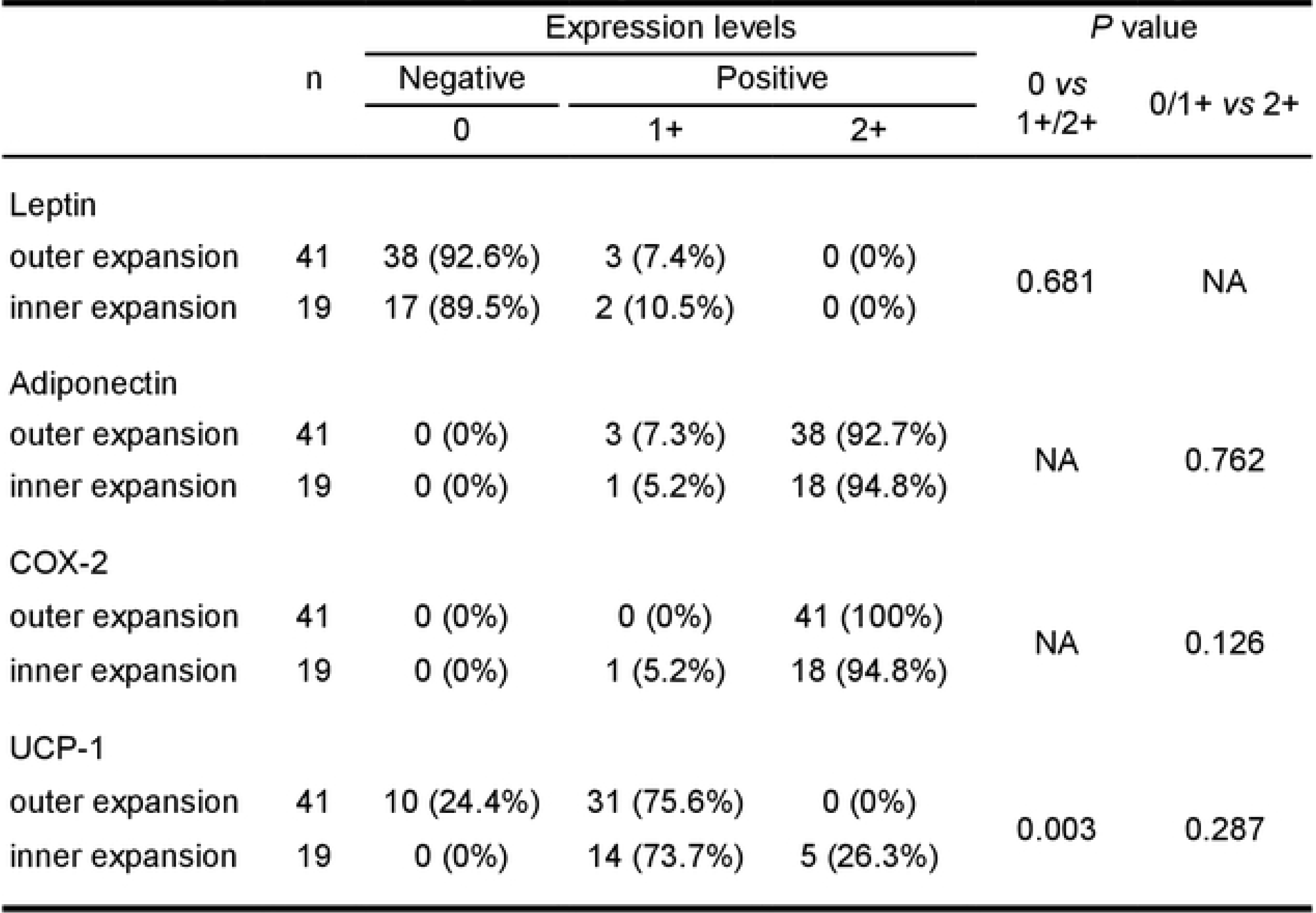
Expression of Leptin, Adiponectin, COX-2 and UCP-1 in perirenal fat.

|  |  | Expression levels |  |  | *P* value |  |
| --- | --- | --- | --- | --- | --- | --- |
|  | n | Negative | Positive |  | 0 vs<br>1+/2+ | 0/1+ vs 2+ |
|  |  | 0 | 1+ | 2+ |  |  |
| Leptin |  |  |  |  |  |  |
| outer expansion | 41 | 38 (92.6%) | 3 (7.4%) | 0 (0%) | 0.681 | NA |
| inner expansion | 19 | 17 (89.5%) | 2 (10.5%) | 0 (0%) |  |  |
| Adiponectin |  |  |  |  |  |  |
| outer expansion | 41 | 0 (0%) | 3 (7.3%) | 38 (92.7%) | NA | 0.762 |
| inner expansion | 19 | 0 (0%) | 1 (5.2%) | 18 (94.8%) |  |  |
| COX-2 |  |  |  |  |  |  |
| outer expansion | 41 | 0 (0%) | 0 (0%) | 41 (100%) | NA | 0.126 |
| inner expansion | 19 | 0 (0%) | 1 (5.2%) | 18 (94.8%) |  |  |
| UCP-1 |  |  |  |  |  |  |
| outer expansion | 41 | 10 (24.4%) | 31 (75.6%) | 0 (0%) | 0.003 | 0.287 |
| inner expansion | 19 | 0 (0%) | 14 (73.7%) | 5 (26.3%) |  |  |

**Figure 2.**
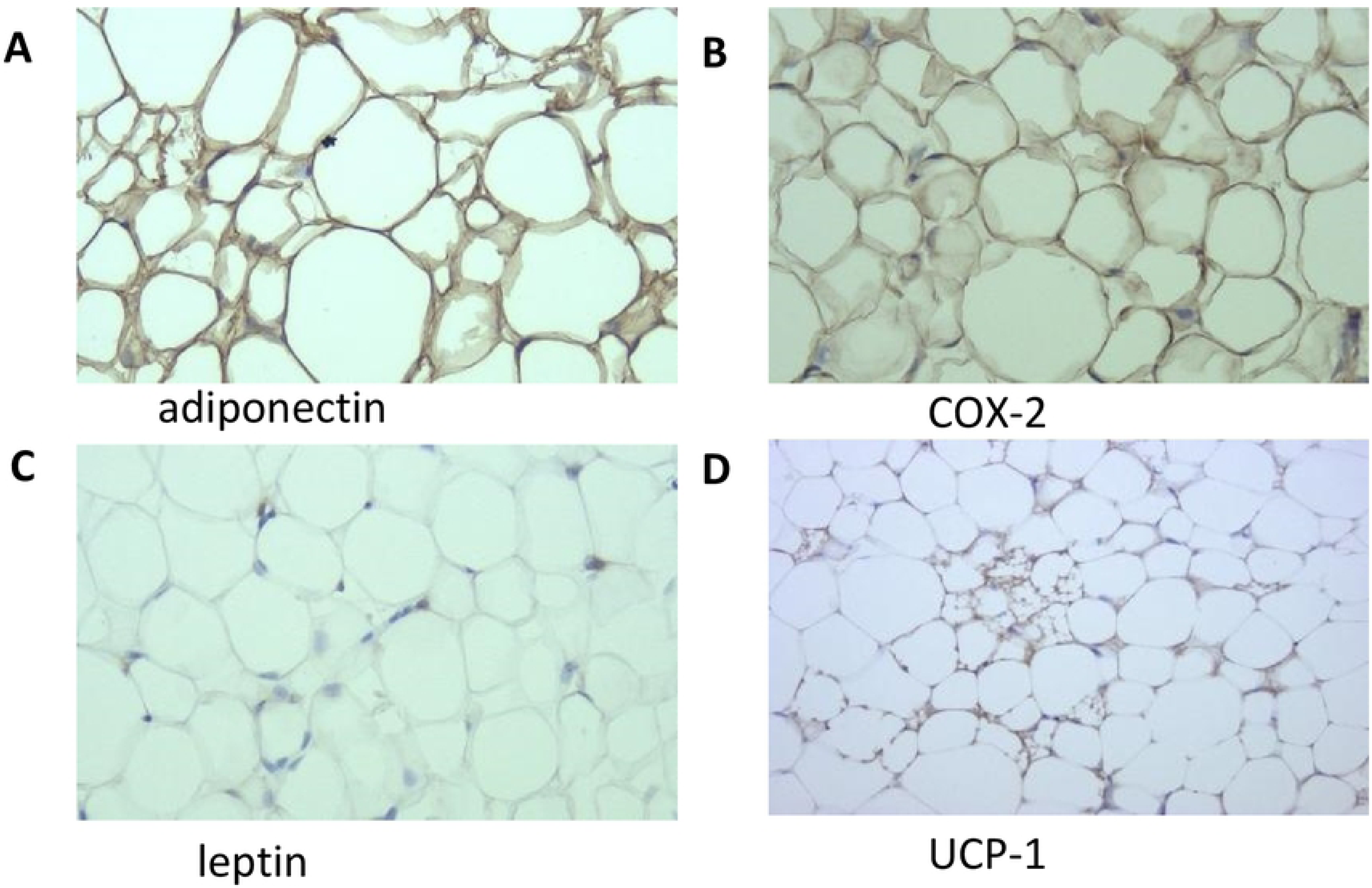
Immunohistochemistry of adiponectin (A), COX-2 (B), leptin (C) and UCP-1 (D) in perirenal fat (original magnification: 200×).

## Discussion

The aim of this study was to determine whether body fat affect the pattern of RCC expansion. Our starting hypothesis was that visceral/perirenal fat affects the growth direction of RCC. Our results support this hypothesis and suggest that perirenal fat plays an important role in the RCC growth pattern.

Around 80% of all body fat is located subcutaneously and 20% is located in visceral areas [17]. The area of visceral fat increases with age, and accumulation of visceral fat increases the risk of not only metabolic diseases [18], but also PCa in men [19]. The V/S ratio is useful for classifying obesity into subgroups: generally, a V/S ratio of 0.4 or above is considered as visceral obesity, and a V/S ratio below 0.4 is considered as subcutaneous obesity [20, 21].

Table 2 showed that pT stage and PF percentage were also independent factors for tumor location. When the tumor becomes larger, it is difficult to stay in the kidney therefore it is reasonable that pT stage is one of the risk factor. Even in pT1a tumor, less than 4cm, PF percentage was correlated with growth pattern (Table S2). Together, these results suggest that perirenal fat can affect the growth direction of renal tumors even if pT stage is the important factor.

In this study, a 50% or more exophytic mass was defined as ‘outer expansion’ because a high level of outgrowth is often easy to resect. Our classification also corresponds with the R.E.N.A.L nephrectomy score, a classification system based on RCC anatomy [22].

Anatomically, kidney cancer can grow into the renal parenchyma or into the perirenal fat. The pressure inside the perirenal fat may be lower than that inside the renal parenchyma; therefore, it may be easier for tumors to grow outside the kidney. This easier growth may also influence the RCC shape and give rise to asymmetrical and/or non-cubic tumors. However, most masses are symmetrical and cubic [22], and it has been reported that RCCs with regular shapes localize to the outside of the kidney and grow more slowly [2]. These studies suggest that the pressure around the RCC may not have a large influence on the direction of growth. We found that perirenal fat may stimulate the RCC to grow into the perirenal fat. Adipose tissue is recognized as the endocrine organ and secrets not only adipocytokines but also biologically effective molecules such as vascular endothelial growth factor, interleukin 6 (IL 6), and TNFα [23]. IL6 and TNFα can induce inflammation and tumorigenesis [24, 25]. However, perirenal adipose tissue secrets certainly these molecules are not definite in this study. Therefore, how the perirenal fat induces the progression and/or carcinogenesis of RCC is unclear. Further studies are required to explore the precise molecular mechanisms involved.

Correlations between adipocytokines, such as leptin and adiponectin, and the clinical characteristics of RCC have been studied. The serum leptin concentration is associated with RCC progression and invasion [26]. In contrast, serum adiponectin is inversely associated with the incidence of RCC [27], and reduced serum adiponectin levels are correlated with increased tumor size and metastasis [28]. The serum leptin concentration is directly associated with BMI, while the adiponectin level is inversely associated with BMI [10]. Leptin interacts with its receptor (ObR) and activates many signals, including VEGF via hypoxia-inducible factor-1α (HIF-1α) and NF-κB [29], and the janus kinase/signal transducer and activator of transcription 3 (JAK/STAT3) [30]. In contrast, adiponectin deficiency suppresses AMPK activation and, as a result, increases angiogenesis in RCC cell lines [31]. These results suggest that adipocytokines play important roles in RCC and may be useful therapeutic targets.

We immunohistochemically stained leptin, adiponectin, COX-2 and UCP-1 in perirenal fat. The expression patterns of leptin and adiponectin did not correlate with the growth pattern (Table 3), in contrast from the results of previous studies of serum levels. In most of our cases, leptin was negative (55 of 60 cases) and adiponectin was positive (60 of 60 cases), and there were few cases in which expression of both adipocytokines were detected. More sensitive modalities will be required in future to quantify these adipocytokines. We also examined COX-2 because obesity can cause adipose tissue overgrowth and inflammation associated with tumorigenesis [32]. However, COX-2 expression also did not correlate with the growth pattern in our study. UCP-1 is a marker of BAT mitochondria [8]. We found that UCP-1 expression was associated with inner-type growth, suggesting that BAT prevents outwards RCC growth. TNFα, which are also secreted by adipose tissue, inhibits UCP1 expression via extracellular-regulated kinases (ERKs) [33], and inflammation also inhibited UCP1 in mice [34]. These results suggest that low expression of UCP1 may reflect the inflammation around the kidney. Furthermore, PTEN, tumor suppressor gene, affect metabolism and regulate *UCP1* transcription [35], therefore UCP1 also reflect the expression of PTEN and as a result, it may suppress RCC protrusion. Further studies are needed to clarify how UCP1 may influence RCC expansion.

We propose that adipose tissue stimulates renal cancer in two ways: (i) adipocytokines that are generated in visceral and/or perirenal fat reach the kidney through the blood stream; or (ii), adipocytokines directly activate RCC through paracrine mechanisms. In this study, we showed that RCCs in patients with thick perirenal fat tend to grow outwardly, such that the RCCs protrude into areas that are rich in adipocytokines and/or inflammatory cytokines. This suggests that paracrine mechanisms are more important for RCC growth direction.

Although thick perirenal fat appears promising, the sample size is relatively small, and this study gave us further hypothesis. We need to investigate the precise mechanisms of adipocyte related signals and also the correlation with cancer progression.

In Conclusion, an increased perirenal fat percentage predicts the RCC growth pattern. Adipose tissue, especially BAT may play an important role in RCC extension. Further studies are needed to investigate the mechanisms involved.

## Acknowlegments

This work was supported by a Japan Society for the Promotion of Science Early Career Scientists Grant (Number 18K16738). We thank Shelley Robison, PhD, from Edanz Group (www.edanzediting.com/ac) for editing a draft of this manuscript.

## Author contributions

EK led the design, analysis, interpretation of data and writing the manuscript. FK and KI performed immunohistochemistry and TA evaluated them. MS, KM, AT, JI, KT and ME contributed to the interpretation of data and reviewing the manuscript.

---

Table S1 Correlations between expansion pattern and characteristics of patients.

Table S2 Correlations between expansion pattern and characteristics of pT1a patients.

## References

1. Autorino R, Khalifeh A, Laydner H, Samarasekera D, Rizkala E, Eyraud R, et al. Robot-assisted partial nephrectomy (RAPN) for completely endophytic renal masses: a single institution experience. BJU Int. 2014;113(5):762–8.

2. Choi SJ, Kim H-S, Ahn S-J, Park Y, Choi H-Y. Differentiating radiological features of rapid-and slow-growing renal cell carcinoma using multidetector computed tomography. J Comput Assist Tomogr. 2012;36(3):313–8.

3. Calle EE, Kaaks R. Overweight, obesity and cancer: epidemiological evidence and proposed mechanisms. Nat Rev Cancer. 2004;4(8):579–91.

4. Cypess AM, Lehman S, Williams G, Tal I, Rodman D, Goldfine AB et al. Identification and importance of brown adipose tissue in adult humans. N Engl J Med. 2009;360(15):1509–17.

5. Marzetti E, D’Angelo E, Savera G, Leeuwenburgh C, Calvani R. Integrated control of brown adipose tissue. Heart and metab. 2016;69:9–14.

6. Kershaw EE, Flier JS. Adipose tissue as an endocrine organ. J Clin Endocrinol Metab. 2004;89(6):2548–56.

7. Vázquez-Vela MEF, Torres N, Tovar AR. White adipose tissue as endocrine organ and its role in obesity. Arch Med Res. 2008;39(8):715–28.

8. Enerbäck S. Human brown adipose tissue. Cell Metab. 2010;11(4):248–52.

9. Hung C-S, Lee J-K, Yang C-Y, Hsieh H-R, Ma W-Y, Lin M-S et al. Measurement of visceral fat: should we include retroperitoneal fat? PLoS One. 2014;9(11):e112355.

10. Galic S, Oakhill JS, Steinberg GR. Adipose tissue as an endocrine organ. Mol Cell Endocrinol. 2010;316(2):129–39.

11. Van Roermund JG, Hinnen KA, Tolman CJ, Bol GH, Witjes J A, Bosch J et al. Periprostatic fat correlates with tumour aggressiveness in prostate cancer patients. BJU Int. 2011;107(11):1775–9.

12. Shiota M, Yokomizo A, Takeuchi A, Imada K, Kiyoshima K, Inokuchi J et al. The feature of metabolic syndrome is a risk factor for biochemical recurrence after radical prostatectomy. J Surg Oncol. 2014;110(4):476–81.

13. Buschemeyer WC, Freedland SJ. Obesity and prostate cancer: epidemiology and clinical implications. Eur Urol. 2007;52(2):331–43.

14. Ewertz M, Jensen M-B, Gunnarsdóttir KÁ, Højris I, Jakobsen EH, Nielsen D et al. Effect of obesity on prognosis after early-stage breast cancer. J Clin Oncol. 2010;29(1):25–31.

15. Thiel DD, Davidiuk AJ, Meschia C, Serie D, Custer K, Petrou SP et al. Mayo adhesive probability score is associated with localized renal cell carcinoma progression-free survival. Urology. 2016;89:54–62.

16. Kashiwagi E, Ide H, Inoue S, Kawahara T, Zheng Y, Reis LO et al. Androgen receptor activity modulates responses to cisplatin treatment in bladder cancer. Oncotarget. 2016;7(31):49169–79.

17. Wajchenberg BL. Subcutaneous and visceral adipose tissue: their relation to the metabolic syndrome. Endocr Rev. 2000;21(6):697–738.

18. Kim SK, Kim HJ, Hur KY, Choi SH, Ahn CW, Lim SK et al. Visceral fat thickness measured by ultrasonography can estimate not only visceral obesity but also risks of cardiovascular and metabolic diseases. Am J Clin Nutr. 2004;79(4):593–9.

19. Hafe P, Pina F, Pérez A, Tavares M, Barros H. Visceral fat accumulation as a risk factor for prostate cancer. Obesity. 2004;12(12):1930–5.

20. Matsuzawa Y, Nakamura T, Shimomura I, Kotani K. Visceral fat accumulation and cardiovascular disease. Obesity. 1995;3(S5).

21. Docimo S, Lee Y, Chatani P, Rogers AM, Lacqua F. Visceral to subcutaneous fat ratio predicts acuity of diverticulitis. Surg Endosc. 2017;31(7):2808–12.

22. Kutikov A, Uzzo RG. The RENAL nephrometry score: a comprehensive standardized system for quantitating renal tumor size, location and depth. J Urol. 2009;182(3):844–53.

23. Kershaw EE, Flier JS. Adipose tissue as an endocrine organ. J Clin Endocrinol Metab. 2004;89(6):2548–56.

24. Hodge DR, Hurt EM, Farrar WL. The role of IL-6 and STAT3 in inflammation and cancer. Eur J Cancer. 2005;41(16):2502–12.

25. Wu Y-d, Zhou B. TNF-α/NF-кB/Snail pathway in cancer cell migration and invasion. Br J Cancer. 2010;102(4):639–644.

26. Horiguchi A, Sumitomo M, Asakuma J, Asano T, Zheng R, Asano T et al. Increased serum leptin levels and over expression of leptin receptors are associated with the invasion and progression of renal cell carcinoma. J Urol. 2006;176(4):1631–5.

27. Spyridopoulos TN, Petridou ET, Skalkidou A, Dessypris N, Chrousos GP, Mantzoros CS. Low adiponectin levels are associated with renal cell carcinoma: A case - control study. Int J Cancer. 2007;120(7):1573–8.

28. Pinthus JH, Kleinmann N, Tisdale B, Chatterjee S, Lu J-P, Gillis A et al. Lower plasma adiponectin levels are associated with larger tumor size and metastasis in clear-cell carcinoma of the kidney. Eur Urol. 2008;54(4):866–74.

29. Gonzalez-Perez RR, Xu Y, Guo S, Watters A, Zhou W, Leibovich SJ. Leptin upregulates VEGF in breast cancer via canonic and non-canonical signalling pathways and NFкB/HIF-1α activation. Cell Signal. 2010;22(9):1350–62.

30. Li L, Gao Y, Zhang L-L, He D-l. Concomitant activation of the JAK/STAT3 and ERK1/2 signaling is involved in leptin-mediated proliferation of renal cell carcinoma Caki-2 cells. Cancer Biol Ther. 2008;7(11):1787–92.

31. Kleinmann N, Duivenvoorden WC, Hopmans SN, Beatty LK, Qiao S, Gallino D et al. Underactivation of the adiponectin–adiponectin receptor 1 axis in clear cell renal cell carcinoma: implications for progression. Clin Exp Metastasis. 2014;31(2):169–83.

32. Pérez-Hernández AI, Catalán V, Gómez-Ambrosi J, Rodríguez A, Frühbeck G. Mechanisms linking excess adiposity and carcinogenesis promotion. Front Endocrinol (Lausanne). 2014;5(65):1–17.

33. Valladares A, Roncero C, Benito M, Porras A. TNF - α inhibits UCP-1 expression in brown adipocytes via ERKs. FEBS letters. 2001;493(1):6–11.

34. Sakamoto T, Nitta T, Maruno K, Yeh YS, Kuwata H, Tomita K et al. Macrophage infiltration into obese adipose tissues suppresses the induction of UCP1 level in mice. Am J Physiol Endocrinol Metab. 2016;310(8):E676–E87.

35. Ortega-Molina A, Serrano M. PTEN in cancer, metabolism and aging. Trends Endocrinol Metab. 2013;24(4):184–189.

